# Domain expansion and functional diversification in vertebrate reproductive proteins

**DOI:** 10.1101/2022.01.31.478407

**Authors:** Alberto M. Rivera, Damien B. Wilburn, Willie J. Swanson

## Abstract

The rapid evolution of fertilization proteins can result in remarkable diversity in their structure and function. Many of the proteins in vertebrate egg coats contain copies (1-6) of the ZP-N domain. These ZP-N domains can facilitate multiple reproduction functions, including species-specific sperm recognition. We integrated phylogenetics and machine learning to investigate how ZP-N domains diversified in structure and function. The most C-terminal ZP-N domain of each paralog is associated with another domain type (ZP-C) which together form a “ZP module.” All modular ZP-N domains were phylogenetically distinct from non-modular or free ZP-N domains. Machine learning-based classification identified 8 residues that form a stabilizing network in modular ZP-N domains that is absent in free domains. Positive selection was identified in some free ZP-N domains. Our findings suggest that purifying selection has conserved an essential structural core in modular ZP-N domains, while free N-terminal domains have been able to experience functionally diversify.

## Introduction

One of the evolutionary innovations associated with multicellularity is the specialization and organization of cells into tissues and organs. Construction of such multicellular structures requires cells to manipulate their local environments by the secretion of different proteins. These components assemble into extracellular matrices and contribute to tissue functions such as morphogenesis, tissue repair (Clause and Barker 2013), homeostasis, and cell adhesion (Streuli 2016). Such extracellular proteins (e.g. collagen) often contain peptide motifs that allow assembly into fibrils and higher order structures (Brodsky and Persikov 2005). Extracellular matrices may be diversified by the duplication of whole genes and individual protein domains, providing new targets for natural selection to act upon. Vertebrate egg coat proteins present a clear example where a complex history of domain duplication is tied to the evolution of an essential extracellular structure.

Vertebrate oocytes are surrounded by an elevated glycoprotein envelope that provides protection from the environment and regulates the rate of sperm entry (Monne et al. 2008). This structure goes by many names, termed the zona pellucida (ZP) in mammals, the chorion in fishes, and the vitelline membrane in amphibians, reptiles, and birds (Wilburn and Swanson 2018). Named after the mammalian version, all vertebrate egg coat proteins contain a pair of immunoglobulin-like domains, ZP-N and ZP-C, that together form a polymerization unit called a ZP module (Jovine et al. 2002; Wilburn and Swanson 2017; Bokhove and Jovine 2018). The last common ancestor of vertebrates possessed six paralogous genes (*zp1, zp2, zp3, zp4, zpd, zpax*) that have experienced clade-specific birth and death events. Consequently, the egg coat of each major vertebrate class has a different composition of ZP module-containing proteins (Conner et al. 2005; Wong and Wessel 2005; Goudet et al. 2008; Meslin et al. 2012; Shu et al. 2015; Wassarman and Litscher 2016; Killingbeck and Swanson 2018). ZP modules are also found in non-reproductive proteins that form extracellular matrices, such as uromodulin (UMOD) which protects against urinary pathogens (Brunati et al. 2015; Bokhove et al. 2016; Devuyst and Pattaro 2018) and tectorin alpha (TECTA) which function in inner ear organization (Bokhove et al. 2016; Kim et al. 2019).

While both ZP-N and ZP-C are immunoglobulin-like domains with a core β-sandwich (Bokhove and Jovine 2018), they are evolutionarily distinct domains that have low amino acid sequence identity, unique disulfide patterns, and variable loop structures (Lin et al. 2011). Independent ZP-C domains outside of the ZP module have been identified in *C. elegans* (Weadick 2020), and four of the egg coat proteins (ZP1, ZP2, ZP4, and ZPAX) contain additional ZP-N domains independent of the ZP-N/ZP-C pair in the ZP module. We refer to ZP-N domains in the module as “modular” and the N-terminal repeats as “free” domains. As ZP-N domains can form asymmetric dimers through their β-sandwich edges (Jovine et al. 2002; Bokhove and Jovine 2018; Litscher and Wassarman 2020), they have been considered the major driver of ZP module polymerization. While free ZP-N domains may similarly function as polymerization units, recent structural studies support that they may have acquired novel functions: the free ZP-N domains of ZP1 form intermolecular cross-links important for egg coat structure (Nishimura et al. 2019), while N-terminal domains in ZP2 (Avella et al. 2013; Avella et al. 2014) and ZP4 (Dilimulati et al. 2022) have been implicated in sperm-egg binding. Despite their functional significance, the evolutionary history of ZP-N domains within and between these many paralogous proteins has not been examined. Using a combination of phylogenetic and machine learning approaches, this manuscript addresses how a complex history of whole gene and tandem domain duplications followed by structural adaptation produced the current diversity of ZP proteins.

## Results and Discussion

We investigated the evolutionary history of vertebrate ZP-N domains by extracting a total of 1448 ZP-N domain sequences from ZP module containing genes of 210 species with both reproductive (*zp1, zp2, zp3, zp4, zpax, zpd*) and non-reproductive (*umod, tecta, cuzd1*) functions (Table S1). While modular and free ZP-N sequences were found to share little sequence identity beyond 4 conserved cysteine residues that form stabilizing disulfide bonds, both domain types are highly similar in three-dimensional structure (Fig 1A). As such, we used a structure-based sequence alignment (Pei et al. 2008) to perform phylogenetic analysis.

**Figure 1:**
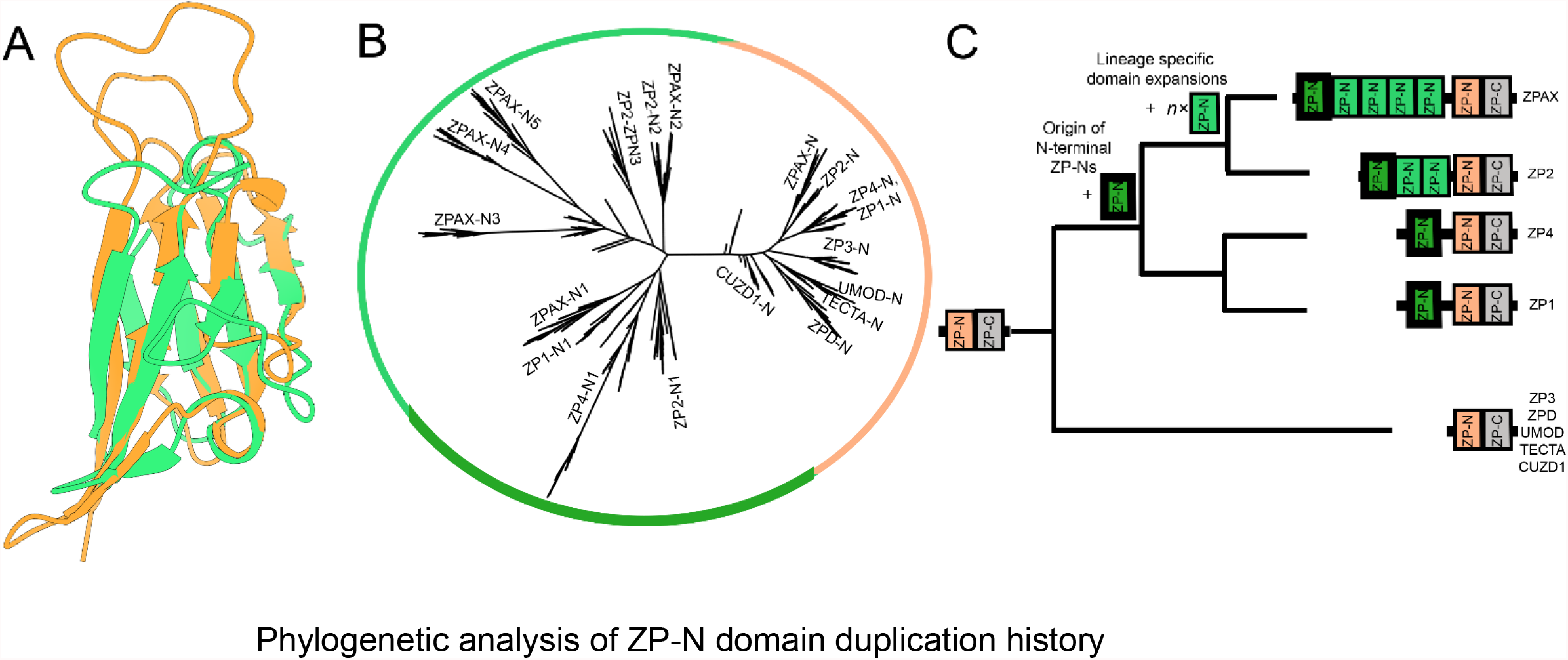
A) A structural alignment of Mouse ZP2-N1 and ZP3-N highlights the broad structural conservation of these two classes of ZP-N domains (RMSD = ∼4.7 Å) despite only ∼18% amino acid sequence identity. B) Phylogenetic analysis (Kozlov et al. 2019) of ZP-N sequences (shown as a maximum likelihood tree) supports an ancestral separation between free and modular ZP-N domains (∼78% support). C) A summary of ZP-N domain evolution based on the gene tree in panel B. The ancestral protein contained a ZP module with a C-terminal ZP-N and ZP-C domains, and duplication of the ZP-N produced the most N-terminal domain found in ZP1, ZP4, ZP2, and ZPAX. Later duplication events within ZP2 and ZPAX gave rise to multiple additional ZP-N domains between ZP-N1 and the ZP module.

Maximum likelihood-based phylogenies indicated that the free ZP-N domains form a single clade distinct from the ZP-C associated modular ZP-N domains (Fig 1B), and this separation was robust to amino acid substitution matrices (LG, WAG, and JTT) (Fig S1). The topology of the modular ZP-N clade was broadly consistent with previously published gene trees based on the complete ZP module with both ZP-N and ZP-C (Claw and Swanson 2012; Feng et al. 2018). The topology of the free ZP-N clade supports that the initial duplication gave rise to the first repeat of the tandem array shared by ZP1, ZP2, ZP4, and ZPAX, which was followed by lineage specific repeat expansions of free ZP-Ns in *ZP2* and *ZPAX* (Fig 1C).

The phylogenetic separation of modular and free ZP-N domains using a structure-based alignment suggests important structural differences between the two domain types, but their high sequence divergence complicated manual identification of such characteristics. Machine learning methods have been applied to various aspects of protein biology such as function prediction (Yang et al. 2018; Bonetta and Valentino 2020) and the classification of membrane bound proteins (Guo et al. 2019). Here, we used a machine learning-based classification strategy to identify what structural features distinguish these two types of ZP-N domains. We applied a logistic regression model to the structurally aligned ZP-N domain sequences, where the probability of being a modular vs free ZP-N type was estimated for each of the 20 amino acids at each position in the alignment. Given the large number of parameters in this model (7981), we combined elastic net regularization and cross-validation to identify the most parsimonious model (i.e. the fewest non-zero parameters) within the 95% confidence interval of the highest scoring model (Fig 2A-B). Through this regularization strategy, we identified a total of 8 modular-associated residues that were sufficient to predict whether a given ZP-N sequence was modular or free with 100% accuracy (Fig 2B).

**Figure 2:**
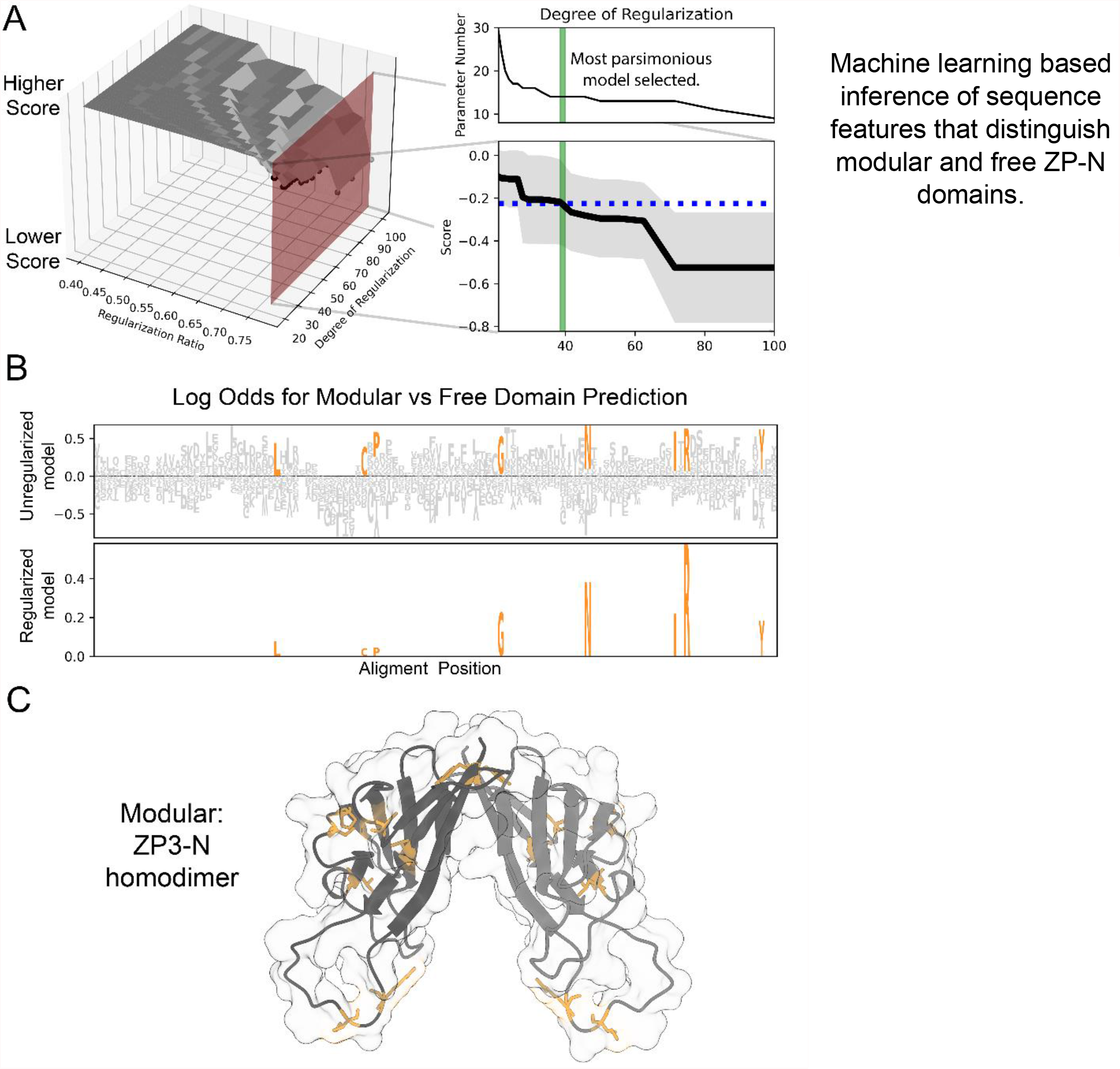
A logistic regression model with elastic net regularization was trained on the ZP-N multiple sequence alignment generated as part of the phylogenetic analysis, with the data partitioned for training and testing (75% and 25%, respectively), with 5-way cross validation of the training data employed to estimate the error distribution of the score function. We defined our optimal model as the most parsimonious model (i.e. the fewest parameters) within the estimated 95% confidence interval of the unregularized model. (A) The space of regularization hyperparameters was explored during model optimization, plotted as a 3D surface (left). The score is the negative mean squared error, and dots correspond to the two-dimensional cross section shown on the right, with the blue line denoting the intersection between the lower confidence limit of the unregularized model to its intersection with the score as a function of regularization strength. B) Comparison of the unregularized and optimal logistic regression models as LOGO plots with the height of each amino acid at each position corresponding to its parameter weight, with colored amino acids denoting parameters retained in the regularized model. Each parameter weight approximating the logs odd ratio for a modular domain prediction, when a residue is present at that position. C) Mapping highly predictive sites onto ZP-N protein models suggest differences in structural properties between free and modular domain. The available crystal structure ZP3-N (3d4c) was used and modelled as a dimer for spatial context. Modular-associated sites are generally buried along the outer edge of the homodimer.

Examination of the residues associated with either ZP-N type in the context of three-dimensional structures suggest differences in both function and quaternary structural dynamics. ZP-N monomers have an immunoglobulin-like β-sandwich fold with the 4- and 3-membered β-strands connected by a disulfide bridge on each edge of the molecule. Biochemical and crystallographic studies support that modular ZP-N domains form asymmetric dimers through the molecular edge that includes the most N- and C-terminal β-strands (Jovine, et al. 2006; Bokhove, et al. 2016). Free ZP-N domains do not appear to dimerize through this N/C-terminal edge, and have experienced functional diversification of the outer edge of the molecule to perform additional protein binding functions (Raj, et al. 2017; Nishimura, et al. 2019). When the modular-associated sites were mapped onto their respective structures, we observed that modular-associated residues form an integrated network of mostly hydrophobic stabilizing contacts that interlock between the β-sheets around the outer edge of the molecules (Fig 2C, Fig S2). The phylogenetic clustering of free ZP-N domains (Fig 1C), along with molecular dynamics support the loss of dimerization activity along the free ZP-N lineage, which could have facilitated their evolution of new binding partners (Fig S3). The stabilizing contacts along the outer edge of the modular ZP-N domains are consistent with these domains principally having structural roles, while in free domains this edge has diversified to allow functional innovation.

Further subdivision of free ZP-N domains by their major clades (the first repeat versus internal repeats in ZP2 and ZPAX) provided little additional information distinguishing these different free ZP-Ns from one another or modular ZP-N domains (Fig S4). Consequently, our sequence-based machine learning classifier appears to have identified residues underlying structural differences between the two domain types that have implications on their respective functions.

The difference in relative conservation of modular domain structures motivated additional analysis of the sequence evolution of these ZP-N domains. Here we focused on mammalian ZP genes (*zp1, zp2, zp3* ,*zp4, umod, tecta*, and *cuzd1*) due to both higher genomic assembly quality and to avoid synonymous substitution saturation that may occur when considering greater phylogenetic breadth (Anisimova and Liberles 2012). Measures of sequence diversity within and between ZP-N groups reveal that modular domains are less diverse overall, and that free ZP-Ns are just as dissimilar to one another as they are to modular domains (Fig 3A).

**Figure 3:**
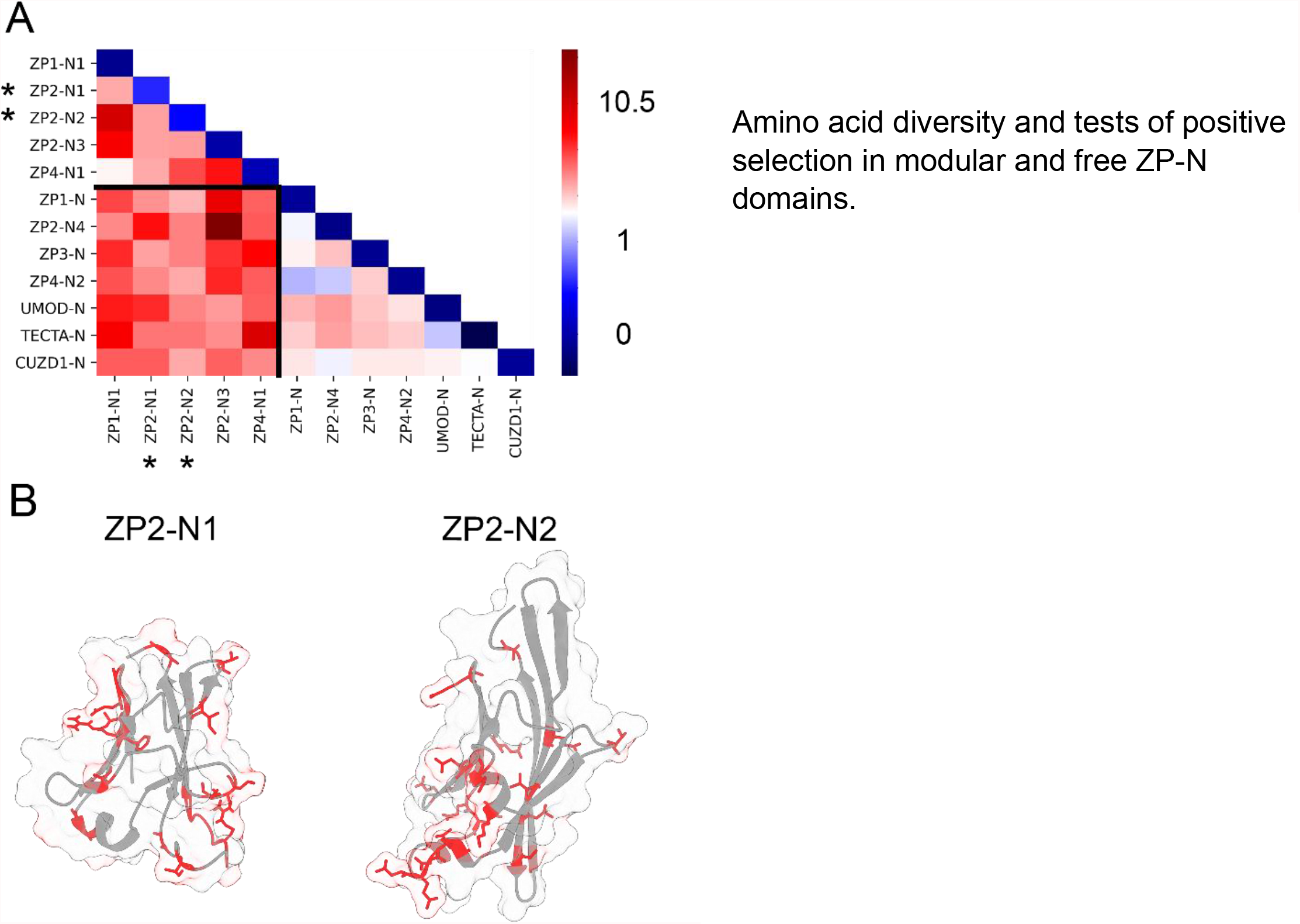
A) A heatmap showing the within group and between group mean phylogenetic distances for orthologous groups of ZP-N domains (Kumar et al. 2018). B) Positively selected sites in mammalian ZP2-N1 and ZP2-N2 were identified through maximum likelihood analysis and mapped onto protein models (4wrn for ZP2-N1, and an AlphaFold prediction for ZP2-N2) (Yang 2007).

These findings motivated molecular evolutionary analyses, and of the 12 ZP-N domains analyzed, only ZP2-N1 and ZP2-N2 showed evidence of positive selection (Table S2). These are notably the two domains with the lowest within group similarity (diagonal of Fig 3A). Positively selected sites in ZP2-N1 are far from the homodimerization edge and physically closer to the network of modular biased residues (Fig 3B). Positively selected sites also constituted a substantial portion of the solvent exposed surface area (34% in ZP2-N1 and 24% in ZP2-N2), potentially facilitating their evolution of novel functions and protein interactions. The rapid evolution of ZP2-N1 is consistent with its role in species-specific sperm recognition (Avella, et al. 2014) and may reflect sexual coevolution with its sperm receptor (whose identity is currently unknown). Remarkably, these positively selected sites cluster near a region associated with species-specific sperm protein binding in free invertebrate ZP-N domains (Raj et al. 2017).

However, based on expansion and retraction of loop lengths outside the core β-sandwich, we believe that these invertebrate free ZP-N domains evolved independently of the free ZP-N domains of vertebrates, suggesting that the expansion of ZP-N arrays for species-specific sperm recognition is a convergent phenomenon that has arisen multiple times throughout metazoan evolution.

In summary, our combined phylogenetic, machine learning classification, and positive selection analyses illustrated a clear distinction between modular and free ZP-N domains. These two classes of domains experienced different evolutionary trajectories, as modular ZP-Ns likely retained a conserved structural role while free ZP-Ns neofunctionalized to serve different reproductive functions. These findings are of relevance to the evolution of species-specificity in fertilization, as the ZP-N domain expansion of ZP2 provided substrates to evolve novel species-specific interactions. Structural changes within free ZP-Ns could result in of a dimerization edge and the evolution of a new sperm binding loop. As these domains are coopted into a reproductive context, coevolution (Clark et al. 2009; Hart et al. 2018) and sexual conflict (Gavrilets and Waxman 2002) with sperm proteins could contribute to their rapid evolution. This reflects the evolutionary dynamics that drive structural diversification and neofunctionalization of duplicated domains. Our combined phylogenetic and machine learning approach outlined here can be applied to other essential gene families with complex duplication histories.

## Materials and Methods

### Multiple Sequence Alignment

Sequences for multiple ZP-N containing proteins were curated from the Ensembl database (release 104) (Howe et al. 2021). Sequences were assigned labelled as one of the ZP genes of interest based on PSI-BLAST e-value scores (Altschul et al. 1997). Sets of orthologous genes were aligned preliminarily with MAFFT (Katoh and Standley 2013) and then trimmed to individual ZP-N domains. Groups of orthologous ZP-N domains were deemed “orthogroups”. Sequences with ambiguous characters were removed, and then sets of orthologous ZP-N sequences were realigned with MAFFT. A full multiple sequence alignment was made by putting the individual orthogroup alignments together using a representative paralog alignment. Individual representative sequences were taken from each orthgroup, and these paralogs were aligned using the structural based PROMALS tool (Pei et al. 2008). This approach was used because of the low sequence identity, but high structural similarity between paralogous Z-N domains. A custom script was used to algorithmically add gaps to orthogroup alignments to form a full multiple sequence alignment. CD-Hit was used to remove highly cluster highly similar sequences (>90% identity) (Li and Godzik 2006; Fu et al. 2012), in order to improve computing speed, and because this study was not concerned with very recent phylogenetic splits.

### Phylogenetics

Maximum likelihood phylogenies were built using RAxML-NG(Kozlov et al. 2019), and multiple different amino acid substitution matrices were tested (LG+G, JTT+G,WAG+G), to demonstrate the robusticity of the deepest phylogenetic divide. The best tree was selected from 100 replications of the maximum likelihood analyses. Nodal support was calculated with transfer bootstrap expectation (Lemoine et al. 2018), a modified form of bootstrapping that is more effective at detecting deep phylogenetic relationships in datasets with large number of taxa. While initial labelling of sequences was based on BLAST values, but when these labels were ambiguous labelling was based on phylogenetic clustering. The clades of ZP1-N1, ZP2-N1, and ZP4-N1, were labelled according to a 90% ma

### Machine Learning

A basic machine learning algorithm using mean squared regression and regularization was coded on python to distinguish the two free and modular groups of ZP-N domains. Logistic regression models are well suited for these classifications, because their outputs are bounded between 0 and 1, which can be interpreted as probability that a given domain is modular(Bewick et al. 2005). The multiple sequence alignment was identical to that used for phylogenetic analysis. The alignment was split into a testing (25%) and training set (75%), and we employed logistic regression modelling with cross-validation on the training set. In that approach, the training set was split into five subsets, and then each of the five subsets is used to validate models trained on the other rest of the training data. For additional rigor, the separate testing dataset was used for the final scoring and model selection.

In order to encode an aligned ZP-N domain within this machine learning framework, each position in the sequence was converted into a vector of twenty digits, corresponding to the twenty amino acids. The value was set to 1 for the entry in the vector corresponding to that residue, and all other values are set to 0. Gapped sites were set to a vector of twenty 0’s. Thus, the classifier was trained using 20n features (plus an additional intercept term), where n is the alignment length. Each of these features has a parameter associated with it and the value of the parameter indicates how informative that feature is, and whether it supports a modular ZP-N or free ZP-N classification. There are a large number of possible parameters in this model (7981 including the intercept), but it is worth considering the optimal number of non-zero parameters for this model. Increasing the number of non-zero parameters will always improve its accuracy on the data it was trained on, but too many parameters will reduce its accuracy when applied to other datasets, in a phenomenon called “overfitting.”(Hawkins 2004)

In optimizing these models, we employed elastic net regularization which two functions that penalize parameters and reduce risk of overfitting (Zou and Hastie 2005). In our sci-kit learn implementation (Pedregosa et al. 2011), we varied both the strength of regularization and the ratio between the two penalty types, and evaluated a range of possible models. The highest scoring model was identified according to the negative mean-squared error scoring metric. In order to choose a suitable sparse model (i.e. fewest non-zero parameters), we adapted the one standard error rule common in machine learning (Hastie et al. 2009), in which you select the sparsest model that is still within one standard error of the highest scoring model. For this analysis we used 95% confidence intervals (∼1.96 standard errors), to achieve the sparsest model (fewest non-zero parameters) that is not statistically different from the highest scoring model sampled. Raw parameter values were plotted in the style of sequence LOGO plots (Schneider and Stephens 1990). The sum of the raw parameter values for matching amino acids in the alignment (and the intercept term), are essentially the log odds that a given sequence is classified as modular. For simplicity, each parameter is described as the log odds associated with a particular residue. After the phylogenetic analysis separated the first N-terminal ZP-N as its own clade, we re-ran the analysis with three multiclassification, but with an otherwise identical hyperparameter search space and computational pipeline.

### Sequence Divergence and Positive Selection Analyses

Our analyses of sequence divergence and positive selection was limited to boreutherian mammals, because of genome quality concerns, and because at large phylogenetic distances sequence differences begin to approach saturation which obscures evolutionary differences (Anisimova and Liberles 2012). For this reason, we were limited to the 12 mammalian ZP-N domains coming from *zp1, zp2, zp3, zp4, umod, tecta*, and *cuzd1*. Boreoeutherian sequences were mined from ensemble (Howe et al. 2021), and were included in these analyses if they were present in 10 or more of these ZP-N domain orthogroups. Phylogenetic distances both within and between orthogroups were calculated in MEGA using poisson estimation with a gamma distribution of variation between sites (Kumar et al. 2016; Kumar et al. 2018).

To determine whether there was any evidence of positive selection we ran PAML analyses (Yang et al. 2005; Yang 2007) on the same sets of ZP-N domains from the sequence divergence estimation. A likelihood ratio test between a model allowing positive selection (M8) and a neutral model (M8a), was used to determine which domains showed evidence of positive selection. Twice the difference of log-likelihood between the models was applied to a chi-squared distribution with one degree of freedom, because M8 has one more parameter M8a. We also performed a Benjamini-Hochberg p-value correction to account for multiple testing (Benjamini and Hochberg 1995). Positively selected sites were visualized on a published crystal structure (ZP2-N1) (Raj et al. 2017), or the alpha-fold predicted structure (Jumper et al. 2021 Jul 15) when this did not exist (ZP2-N2). Sites were labelled if they had a posterior probability of being positively selected > 75% according Bayes Empirical Bayesian (BEB) analysis.

### Visualization and Other methods

When protein structures were not available Alpha-Fold2 tertiary structure prediction was used (Jumper et al. 2021), and three-dimensional protein structures were visualized using either pymol (Schrödinger 2015) or ChimeraX (Pettersen et al. 2004). Rosetta (Chaudhury and Gray 2008; Sircar et al. 2010) was used to perform docking simulations on ZP2-N1 and ZP3-N homodimers, to illustrate the greater energetic stability of modular ZP-N homodimers.

---

**Table S1:**
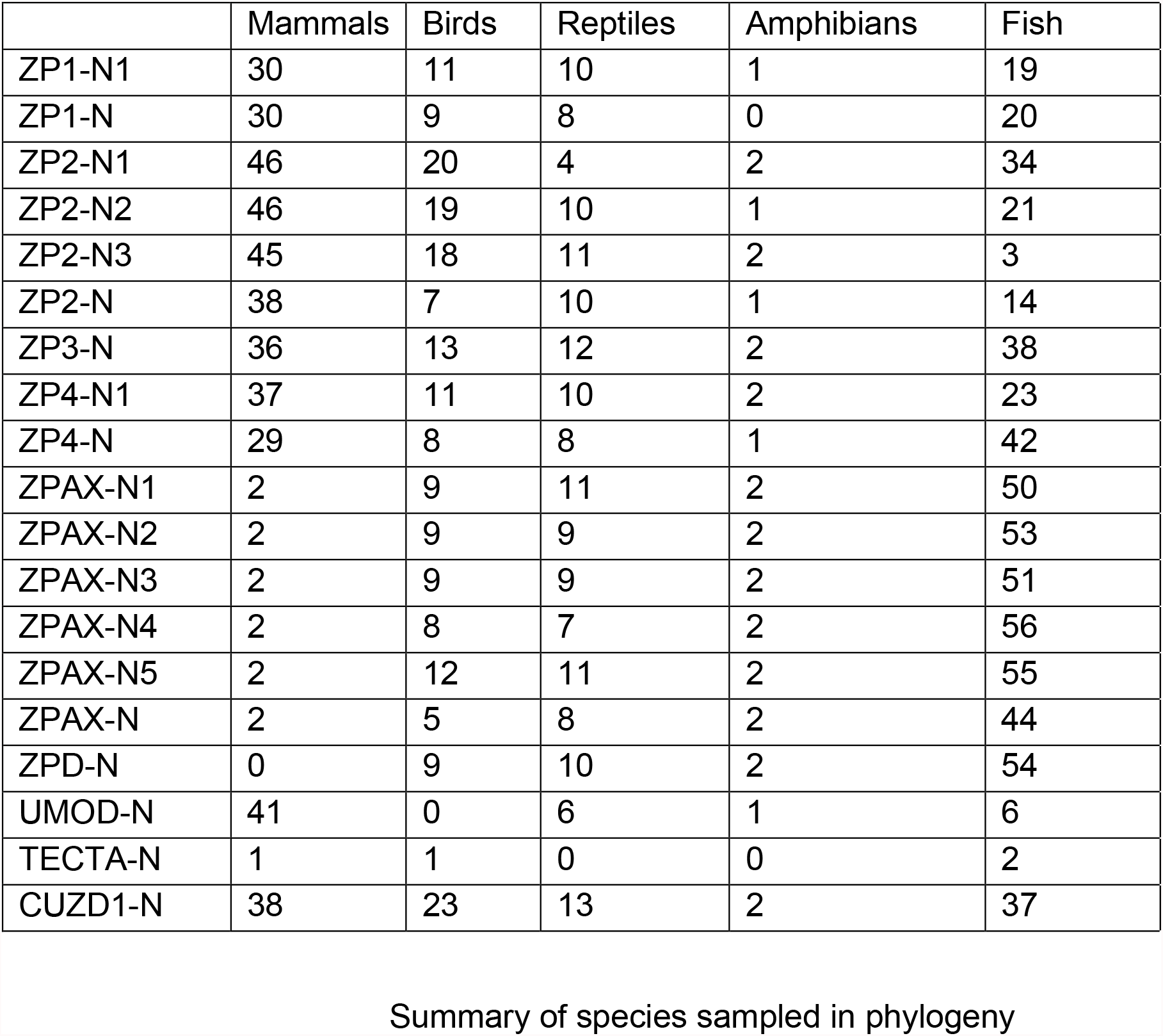
We included a total of 1488 ZP-N domain sequences across the five classes of vertebrates in the final phylogeny. Here the amphibians only included frogs due to genomic availability reasons. The fish class includes all non-tetrapod vertebrates, and is non-monophyletic. Labels in this table are based on BLAST results.

|  | Mammals | Birds | Reptiles | Amphibians | Fish |
| --- | --- | --- | --- | --- | --- |
| ZP1-N1 | 30 | 11 | 10 | 1 | 19 |
| ZP1-N | 30 | 9 | 8 | 0 | 20 |
| ZP2-N1 | 46 | 20 | 4 | 2 | 34 |
| ZP2-N2 | 46 | 19 | 10 | 1 | 21 |
| ZP2-N3 | 45 | 18 | 11 | 2 | 3 |
| ZP2-N | 38 | 7 | 10 | 1 | 14 |
| ZP3-N | 36 | 13 | 12 | 2 | 38 |
| ZP4-N1 | 37 | 11 | 10 | 2 | 23 |
| ZP4-N | 29 | 8 | 8 | 1 | 42 |
| ZPAX-N1 | 2 | 9 | 11 | 2 | 50 |
| ZPAX-N2 | 2 | 9 | 9 | 2 | 53 |
| ZPAX-N3 | 2 | 9 | 9 | 2 | 51 |
| ZPAX-N4 | 2 | 8 | 7 | 2 | 56 |
| ZPAX-N5 | 2 | 12 | 11 | 2 | 55 |
| ZPAX-N | 2 | 5 | 8 | 2 | 44 |
| ZPD-N | 0 | 9 | 10 | 2 | 54 |
| UMOD-N | 41 | 0 | 6 | 1 | 6 |
| TECTA-N | 1 | 1 | 0 | 0 | 2 |
| CUZD1-N | 38 | 23 | 13 | 2 | 37 |

**Table S2:**
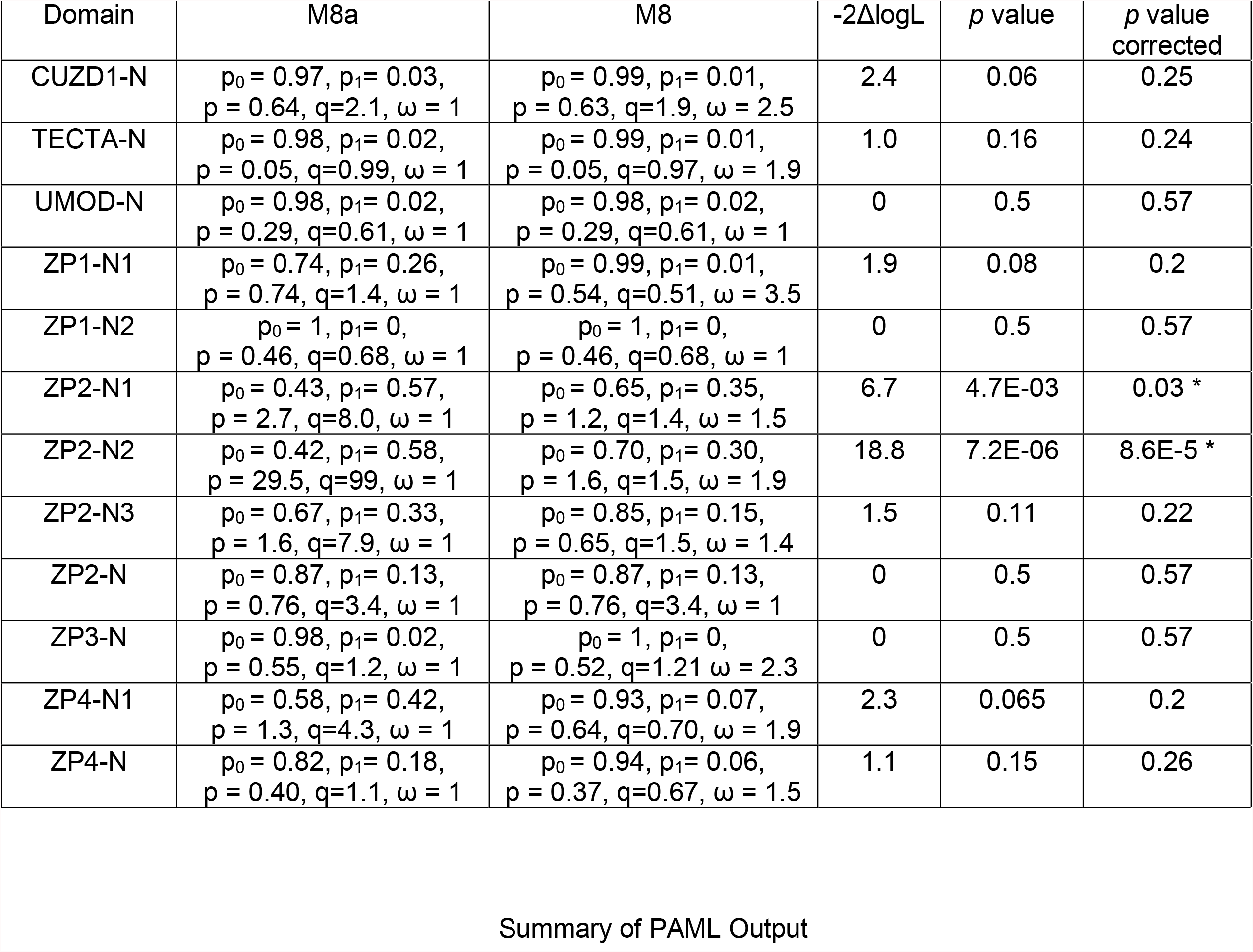
This table summarizes the results from PAML analysis (Yang 2007). Here we compared a neutral model (M8a) to a model that allows positive selection (M8). A * denotes statistically significant p values after Benjamini-Hochberg multiple testing correction (Benjamini and Hochberg 1995).

| Domain | M8a | M8 | -2ΔlogL | *p* value | *p* value corrected |
| --- | --- | --- | --- | --- | --- |
| CUZD1-N | $p_0 = 0.97, p_1 = 0.03,$<br>$p = 0.64, q=2.1, \omega = 1$ | $p_0 = 0.99, p_1 = 0.01,$<br>$p = 0.63, q=1.9, \omega = 2.5$ | 2.4 | 0.06 | 0.25 |
| TECTA-N | $p_0 = 0.98, p_1 = 0.02,$<br>$p = 0.05, q=0.99, \omega = 1$ | $p_0 = 0.99, p_1 = 0.01,$<br>$p = 0.05, q=0.97, \omega = 1.9$ | 1.0 | 0.16 | 0.24 |
| UMOD-N | $p_0 = 0.98, p_1 = 0.02,$<br>$p = 0.29, q=0.61, \omega = 1$ | $p_0 = 0.98, p_1 = 0.02,$<br>$p = 0.29, q=0.61, \omega = 1$ | 0 | 0.5 | 0.57 |
| ZP1-N1 | $p_0 = 0.74, p_1 = 0.26,$<br>$p = 0.74, q=1.4, \omega = 1$ | $p_0 = 0.99, p_1 = 0.01,$<br>$p = 0.54, q=0.51, \omega = 3.5$ | 1.9 | 0.08 | 0.2 |
| ZP1-N2 | $p_0 = 1, p_1 = 0,$<br>$p = 0.46, q=0.68, \omega = 1$ | $p_0 = 1, p_1 = 0,$<br>$p = 0.46, q=0.68, \omega = 1$ | 0 | 0.5 | 0.57 |
| ZP2-N1 | $p_0 = 0.43, p_1 = 0.57,$<br>$p = 2.7, q=8.0, \omega = 1$ | $p_0 = 0.65, p_1 = 0.35,$<br>$p = 1.2, q=1.4, \omega = 1.5$ | 6.7 | 4.7E-03 | 0.03 * |
| ZP2-N2 | $p_0 = 0.42, p_1 = 0.58,$<br>$p = 29.5, q=99, \omega = 1$ | $p_0 = 0.70, p_1 = 0.30,$<br>$p = 1.6, q=1.5, \omega = 1.9$ | 18.8 | 7.2E-06 | 8.6E-5 * |
| ZP2-N3 | $p_0 = 0.67, p_1 = 0.33,$<br>$p = 1.6, q=7.9, \omega = 1$ | $p_0 = 0.85, p_1 = 0.15,$<br>$p = 0.65, q=1.5, \omega = 1.4$ | 1.5 | 0.11 | 0.22 |
| ZP2-N | $p_0 = 0.87, p_1 = 0.13,$<br>$p = 0.76, q=3.4, \omega = 1$ | $p_0 = 0.87, p_1 = 0.13,$<br>$p = 0.76, q=3.4, \omega = 1$ | 0 | 0.5 | 0.57 |
| ZP3-N | $p_0 = 0.98, p_1 = 0.02,$<br>$p = 0.55, q=1.2, \omega = 1$ | $p_0 = 1, p_1 = 0,$<br>$p = 0.52, q=1.21, \omega = 2.3$ | 0 | 0.5 | 0.57 |
| ZP4-N1 | $p_0 = 0.58, p_1 = 0.42,$<br>$p = 1.3, q=4.3, \omega = 1$ | $p_0 = 0.93, p_1 = 0.07,$<br>$p = 0.64, q=0.70, \omega = 1.9$ | 2.3 | 0.065 | 0.2 |
| ZP4-N | $p_0 = 0.82, p_1 = 0.18,$<br>$p = 0.40, q=1.1, \omega = 1$ | $p_0 = 0.94, p_1 = 0.06,$<br>$p = 0.37, q=0.67, \omega = 1.5$ | 1.1 | 0.15 | 0.26 |

**Supplemental Figure 1:**
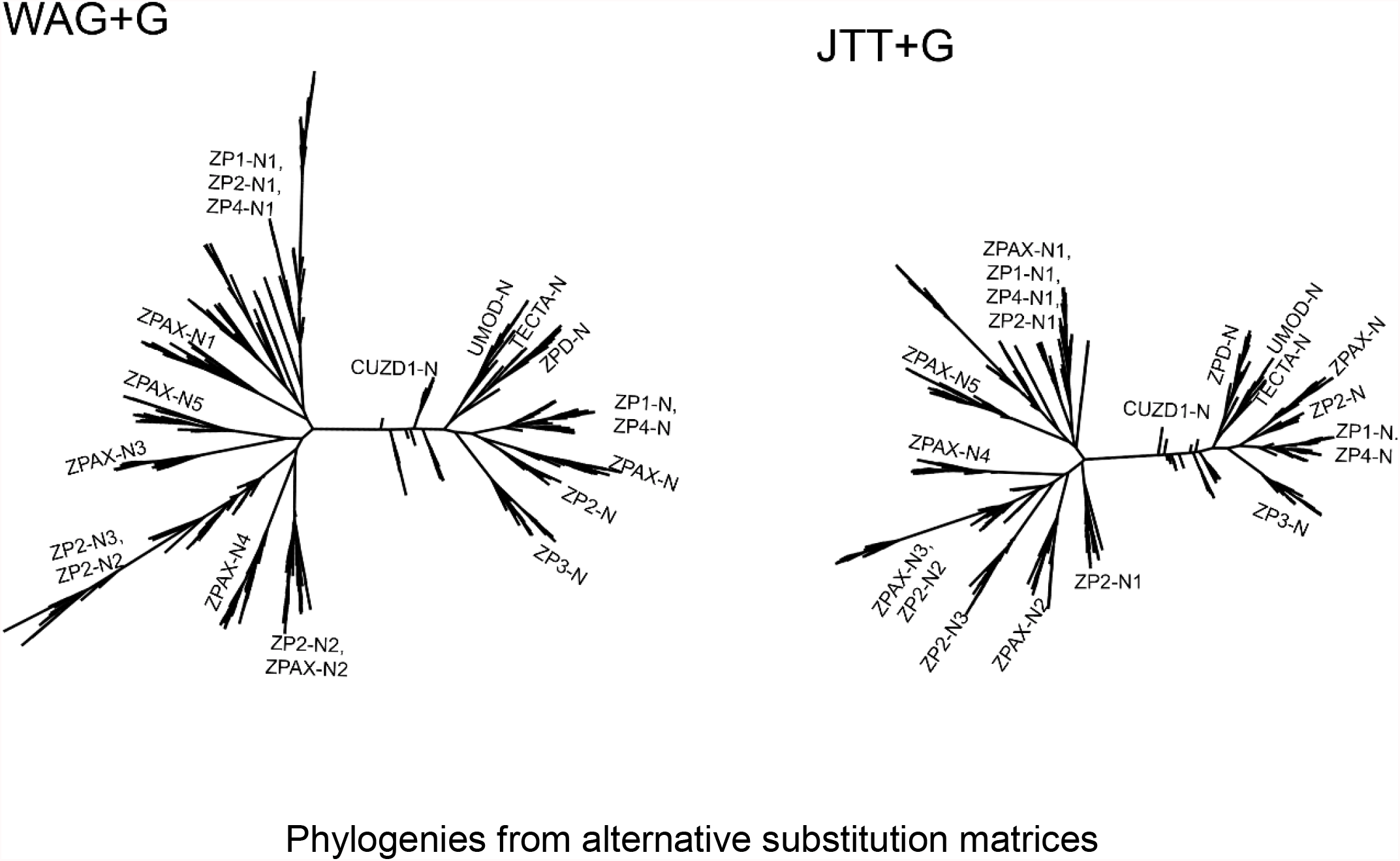
These two trees were produced through RAxML-NG(Kozlov et al. 2019). Major aspects of the tree are conserved, specifically the monophyletic grouping of free ZP-N domains.

**Supplemental Figure 2:**
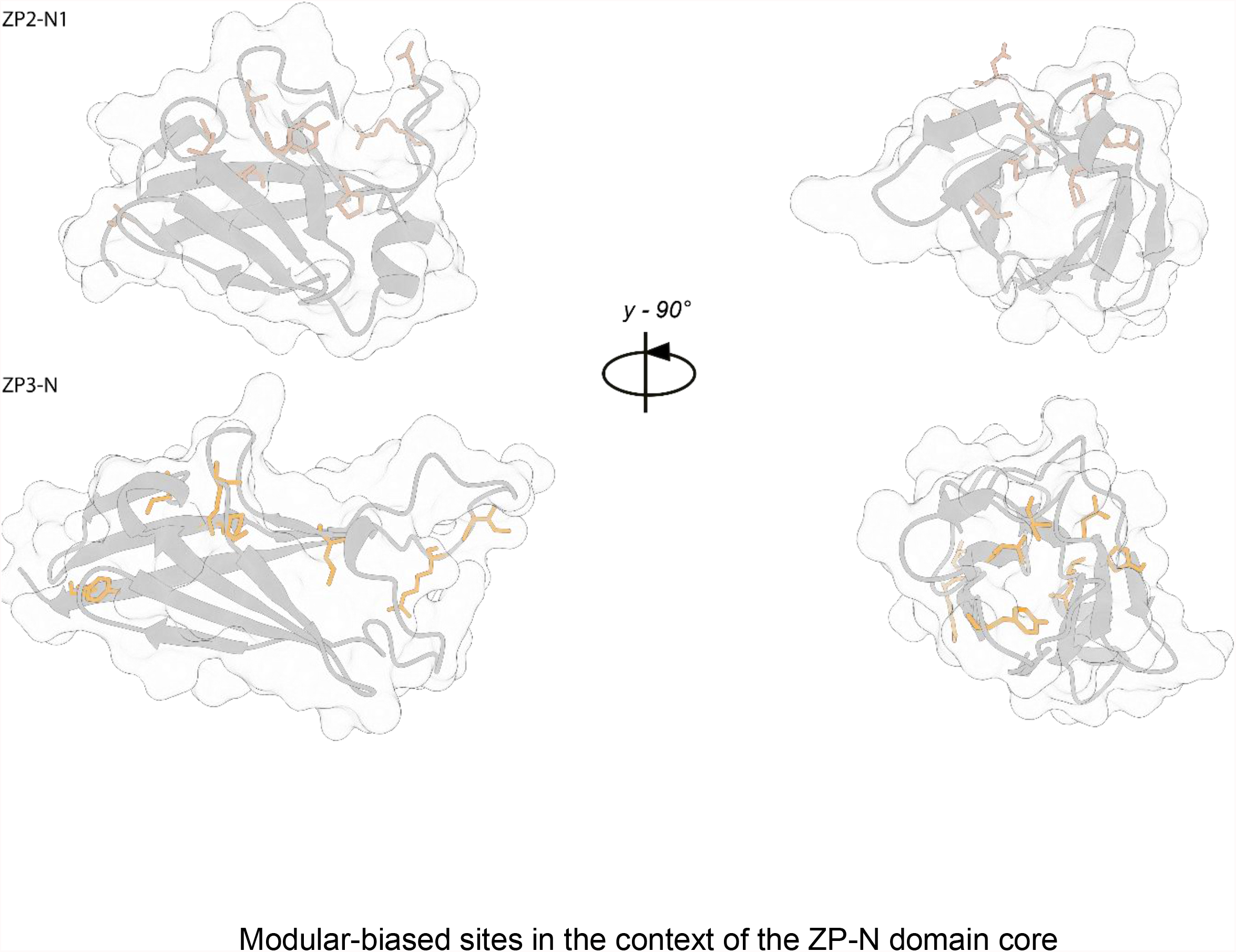
This figure shows the modular biased sites in the context of both ZP2-N1 (Raj et al. 2017) and ZP3-N (Monne et al. 2008) crystal structures. The brighter orange corresponds to the modular-biased sites according to our machine learning model, while the duller orange on ZP2-N1 represent the structural homolog of those sites according to a pymol (Schrödinger 2015) structural alignment. We observe a clustering of these sites within the core of the domain in ZP3-N and not ZP2-N1. The two positions of each are 90° rotations around the y-axis.

**Supplemental Figure 3:**
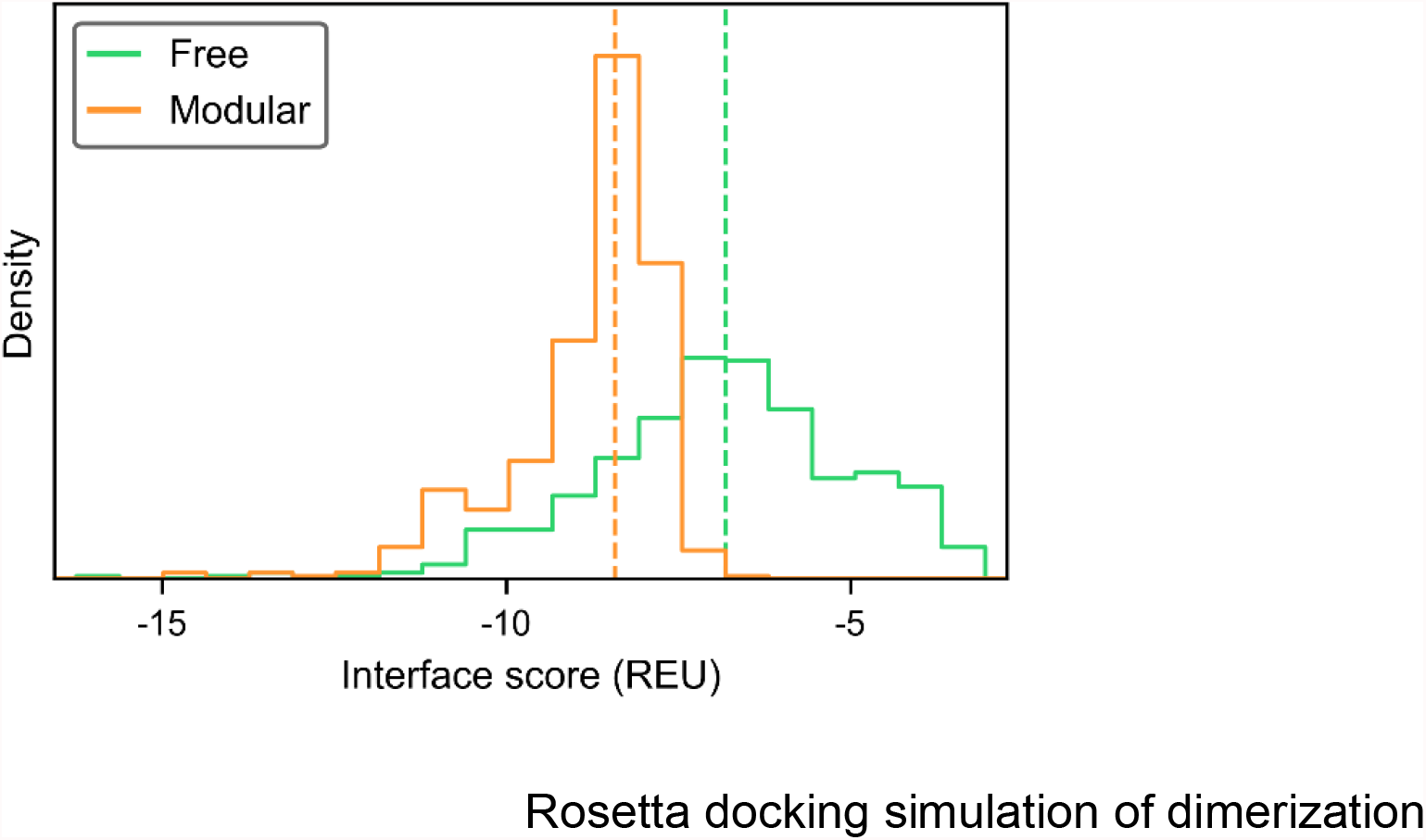
This is a histogram of interface scores from rosetta docking simulations of dimers for both free (ZP2-N1) and modular (ZP3-N) ZP-N domains.

**Supplemental Figure 4:**
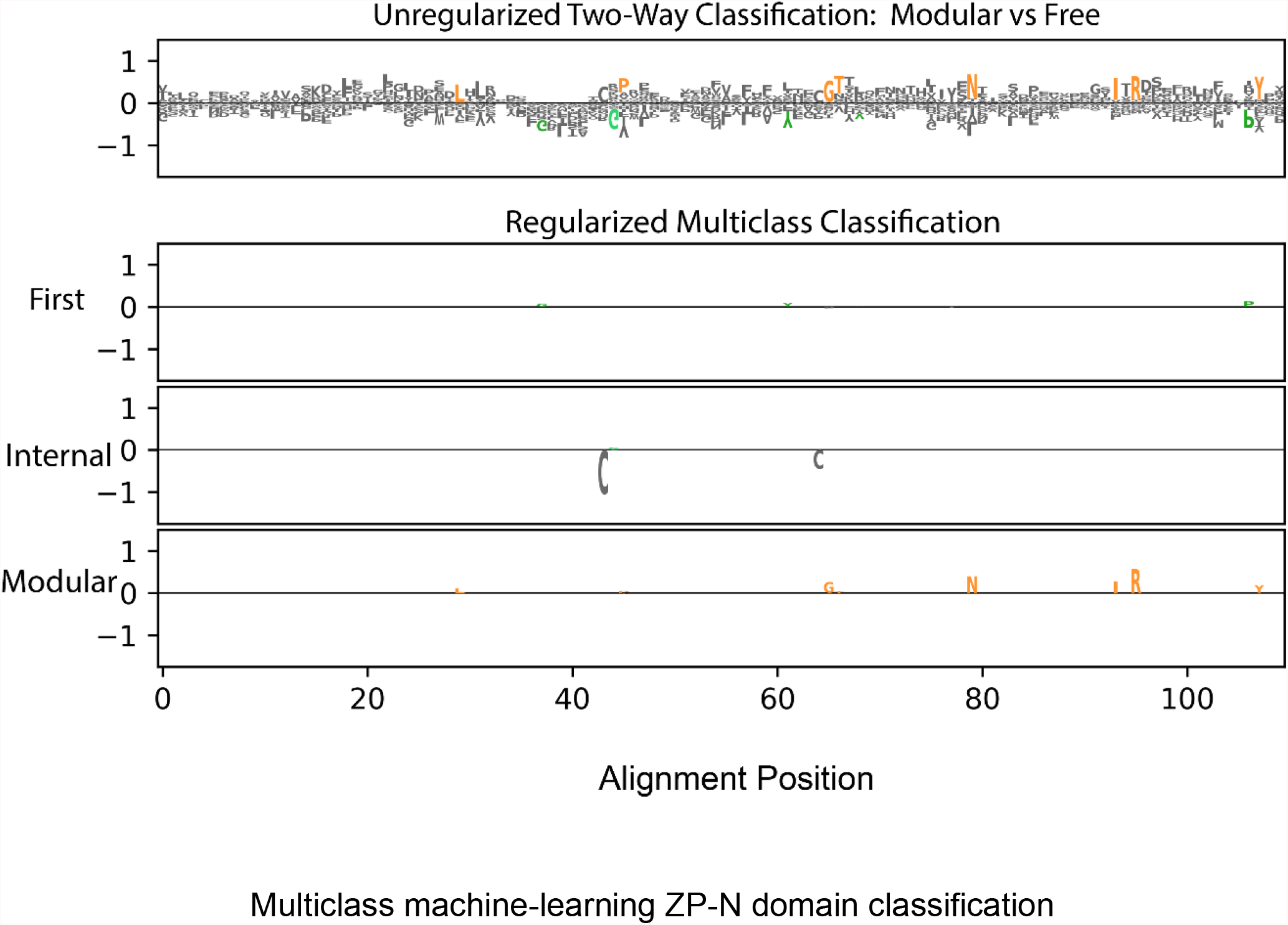
Since we observed monophyletic grouping of the most N-terminal ZP-N domain, we performed a multiclass variation of our machine learning analysis. The data was split into three classes: modular, first (i.e. most N-terminal free domain), and internal domains. In this multiclass analysis the model is fit three times, each time producing a classifier that distinguishes one of the classes from the rest of the data. The first row is the unregularized model from our two-class analysis, and our regularized multiclass models are summarized in the other three rows, where positive values suggest a bias towards that class. Our modular classifier recapitulates our earlier results, because it is in essence still comparing modular ZP-Ns versus all free ZP-Ns. The first ZP-N domain has few amino acids associated with it with relatively low parameter values. The internal ZP-N domains seem to have a bias against the cys 2-3 bond, but that likely is a reflection of the conservation of the second cysteine in the first ZP-Ns and the third cysteine in modular domains

